# Feature selectivity explains mismatch signals in mouse visual cortex

**DOI:** 10.1101/2021.04.12.439457

**Authors:** Tomaso Muzzu, Aman B. Saleem

## Abstract

Sensory experience is often dependent on one’s own actions, including self-motion. Theories of predictive coding postulate that actions are regulated by calculating prediction error, which is the difference between sensory experience and expectation based on self-generated actions. Signals consistent with prediction error have been reported in mouse visual cortex (V1) when visual flow coupled to running is unexpectedly perturbed. Here, we show that such signals can be elicited by visual stimuli uncoupled with the animal’s running. We recorded the activity of mouse V1 neurons while presenting drifting gratings that unexpectedly stopped. We found strong responses to visual perturbations, which were enhanced during running. If these perturbation responses are signals about sensorimotor mismatch, they should be largest for front-to-back visual flow expected from the animals’ running. Responses, however, did not show a bias for front-to-back visual flow. Instead, perturbation responses were strongest in the preferred orientation of individual neurons and perturbation responsive neurons were more likely to prefer slow visual speeds. Our results therefore indicate that prediction error signals can be explained by the convergence of known motor and sensory signals in visual cortex, providing a purely sensory and motor explanation for purported mismatch signals.

## Introduction

Sensation and action are two intertwined processes that the brain continuously executes and adjusts^1–5^. Theories of predictive coding postulate that sensation is an active process that uses information about one’s own actions to distinguish between self-generated and external sensory stimuli. One feature of such predictive coding is the computation of prediction error – the difference between observed features and those expected based on one’s own actions. Prediction error signals have been shown to be encoded in many neural circuits, most famously in the reward system^6^, and also in motor^7^ and sensory brain regions^8,9^.

Locomotion through a familiar environment can generate predictable sensory experiences, including visual flow, that are in turn important in guiding behaviour. It has been suggested that errors in the prediction of visual flow are encoded as early as in the primary visual cortex^10,11^, based on the observation of large responses to sudden stops of visual flow that was normally coupled to an animal’s running. The same visual perturbation (cessation of visual flow) played back to a stationary animal elicited smaller responses. In agreement with theories of predictive coding, such activity would provide V1 with the ability to encode the error between the actual sensory feedback and the expected one, i.e. a visuomotor mismatch signal^10^. It has been suggested that this visuomotor mismatch is driven by the comparison of excitatory, motor inputs with inhibitory, visual flow inputs^12,13^. However, neurons in V1 have a wide range of visual speed (or temporal frequency) preferences^14–15^, and running both drives V1 activity and modulates responses to visual stimuli^10,17–22^. An alternate and untested hypothesis is that responses to sudden stops of visual flow are simply due to the convergence of motor and visual inputs, and do not arise from the precise coupling between an animals’ actions and the visual stimulus.

## Results

To test how changes to visual flow affected neural activity in the mouse visual cortex we presented gratings drifting at a constant speed (0.04 cycles/°, 3 cycles/s, trial duration = 7.3 s), in the right visual field (covering 120° by 120°, Fig. 1a). Mice were head-restrained and free to run on a polystyrene wheel. On each trial, the contrast of the drifting grating slowly increased from 0 to 0.8, and was then held constant for 1s, before decreasing back to 0 again. We chose this protocol so that mice did not experience sudden, transient changes of stimulus appearance or speed. On the recording day, however, the drifting grating was suddenly stopped for one second in a random 25% of the trials, thus causing an unexpected perturbation of the visual flow (Fig. 1b).

**Fig. 1:**
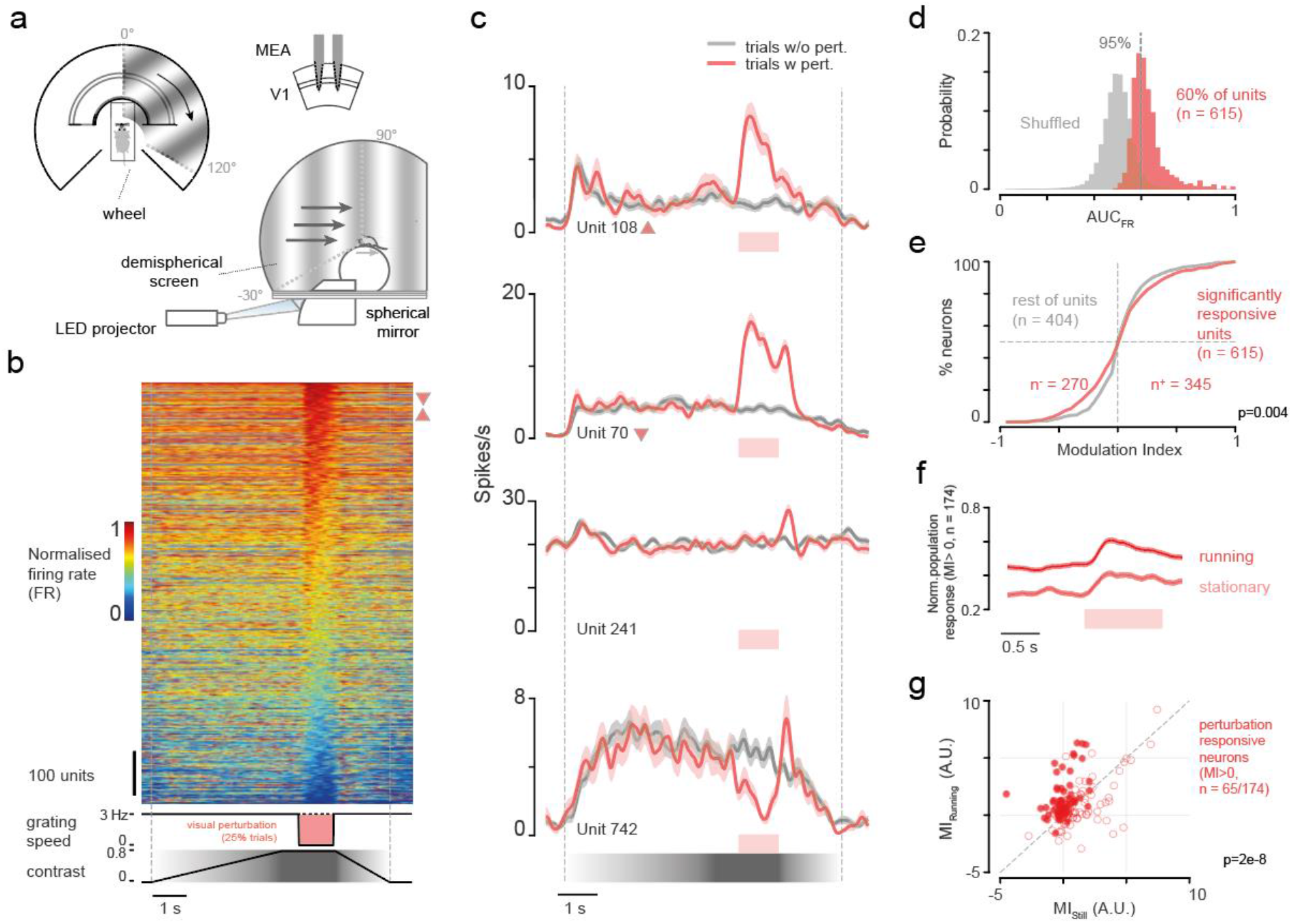
V1 neurons show responses to visual flow perturbations that are stronger during running. **a**, Side and top view of the recording apparatus. Bottom right: schematic of the MEA silicon probe and recording area. **b**, Top: normalised mean response of all recorded units (n=1019) for trials with perturbation. Bottom: typical contrast and temporal frequency of the drifting grating for trials with perturbation (TF=0 cycles/s). Triangles refer to the position of example units shown in c. **c**, Mean response to trials with perturbation (red trace) and without perturbation (black trace) of four example units significantly responsive to perturbation. Unit number is consistent with row in (c). Shaded error bars indicate s.e.m. **d**, Probability distribution of the area under the receiving operating characteristic curve of logistic classifier trained on shuffled and recorded data. Dotted line represents the 95 percentile threshold. e, Probability cumulative distribution of the modulation indexes (MI) of perturbation responsive units (red) and the other units (gray). P value for two-sample Kolmogorov-Smirnov test. **f**, Normalised mean population firing rate activity during perturbation trials in running and stationary conditions. **g**, Modulation of perturbation responses in running trials versus stationary trials for neurons recorded in sessions in which there were at least 4 trials for each condition (n = 174). The modulation of each unit, for each condition, was computed as the sum of all firing rates of the perturbation period minus those evaluated during one second preceding the perturbation. Solid dots show units that were significantly modulated by running individually (non-parametric test with significance threshold at p=0.05, see methods for details).

The perturbation of purely visual flow was clearly reflected in the activity of neurons recorded in V1 (Fig. 1b). We used multi-electrode array silicon probes to record the activity of neurons in the visual cortex. Many individual neurons showed a change in their firing rates during the perturbation (Fig. 1b, c). To quantify the reliability of each neuron’s perturbation response, we trained a logistic regression classifier to discriminate between perturbation and non-perturbation trials based on features of an individual neuron’s activity, including response amplitude during the visual flow halts (Fig. 1c, d, see methods for details). The classifier would only discriminate between the trial types when the perturbation responses are large and reliable. We considered a unit reliable if the classifier performed better than 95% of classifiers trained on shuffled data (Fig. 1d), and found 60% of neurons (n=615/1019) reliably responsive to visual perturbation. As the classifier could use either positive or negative perturbation responses, we also characterised the sign of the responses by measuring a modulation index (MI): the proportional change in firing rate during the perturbation, relative to the pre-perturbation period. We found that most neurons responded to the perturbation by increasing their firing rate (n=345 of 1,019 vs. 270 of 1,019 units that reduced responses, Fig. 1e).

Perturbation responsive neurons appear to respond as if they encode visuomotor mismatch: activity increases when the visual flow, otherwise coupled to the animal’s running, is suddenly stopped. We therefore evaluated the effects of running on the perturbation responses, in sessions that had at least 5 trials in both running (speed ≥2 cm/s) and stationary conditions. While perturbation responses were present in both stationary and running conditions, they were markedly different: pre-perturbation activity was higher during running (Fig. 1f; p=0.002, Wilcoxon sign-rank test), and the modulation index was higher when the animal was running compared to when it was stationary (n=65/174 units, p<10^-7^, Wilcoxon sign rank test). Therefore, the perturbation responses were larger when animals were running.

Visual perturbation responses were not explained by perturbation-induced changes in running behaviour. We quantified changes in running speed in each recording session using the same metrics as for neural responses, calculating reliability and modulation indices for speed changes. Mice showed reliable changes in speed following the perturbation in less than half the sessions (n=19/47 sessions with 95 percentile performance, Suppl. Fig 1a, b), and these sessions had a similar fraction of perturbation units as sessions with no running speed changes (35% compared to 34% in other sessions). Importantly, trial-by-trial modulation indices of neural responses were rarely (n = 8/315 units) correlated with modulation indices of speed changes (Suppl. Fig. 1c, d, e). Therefore, neither the occurrence nor the change in running speed explained perturbation responses.

A crucial feature of the sensorimotor mismatch hypothesis is that perturbation responses will be largest for the motion direction predicted by the animals’ running (front-to-back, or naso-temporal flow). Previous reports of responses during sensorimotor mismatch were measured using stimuli moving only along this direction, leaving open the question of whether perturbation responses might also be present for other motion directions. We therefore tested the perturbation response in eight different directions – in each case the stimulus moved in a particular direction and suddenly stopped during the perturbation period.

We found that population perturbation responses were similar in every motion direction tested (Fig. 2). For each direction, we measured the population perturbation response as the average z-scored response across all the perturbation units within a recording session (n=47 recordings). The population perturbation response did not display a preference for any direction (Suppl. Fig. 3b) or orientation (Fig. 2b).

**Fig. 2:**
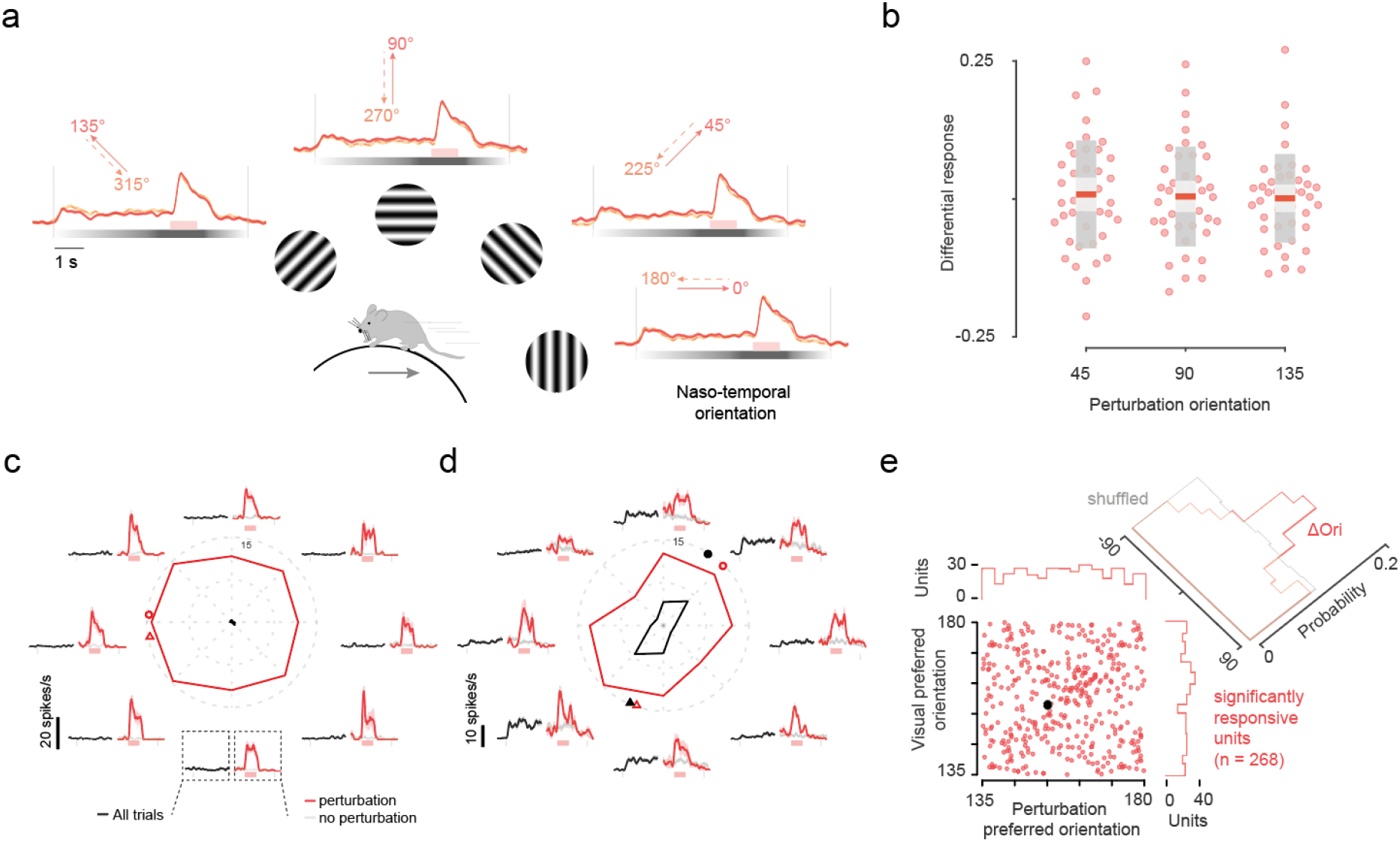
Perturbation responses are not biased to the front-to-back direction and are stronger at the preferred orientation of single neurons. **a**, Mean normalised population responses of the perturbation responsive units (only positively modulated) for different grating directions. Dotted lines indicate the start/end of the trial and of the perturbation period. **b**, Perturbation responses to orientation angles for all recording sessions (n = 41) minus responses to naso-temporal orientation. Colours of box plots indicate mean (red), s.e.m. (light gray), s.t.d. (dark grey). **c,d**, Mean responses of two example units for the different grating directions. Central panel of each example shows the mean responses for different grating directions computed during the first 4 seconds of the trials minus the baseline activity (black line). Red line is the mean firing rate computed during the visual perturbation period (same baseline as the grating stimulus is used to avoid any bias from visual direction/orientation tuning of the unit). Surrounding plots show mean responses for each direction. The responses to the visual stimulus (0 - 4 s) are averaged over all trials in the direction. Responses to the visual perturbation are averaged over the number of trials for each condition (red: with perturbation; light gray: no perturbation). Circles indicate preferred orientation. Triangles indicate preferred direction. **e**, Scatter plot of preferred orientation angle for grating stimulus and visual perturbation. Black data point indicates example unit shown in D. Only units tuned to orientation are orientation are considered (Hotelling’s t-squared test, p<=0.01 n = 268).

We next asked if individual neurons showed any preference to the direction of the perturbation. While some neurons showed no preference for the orientation of the perturbation (Fig 2c), we found some neurons that had a higher response along certain orientations (Fig 2d-e). We defined the orientation with the maximal perturbation response as the preferred perturbation orientation. The distribution of preferred perturbation orientations did not show a bias for any particular orientation or direction (Fig 2e & Suppl. Fig. 3c), which is consistent with the lack of bias in the population responses. Therefore, the perturbation responses were not biased in the front-to-back direction as predicted by a sensorimotor mismatch.

The perturbation responses were influenced by the orientation selectivity of the neuron (Fig. 2c-e). We measured the orientation selectivity in the period before the perturbation onset (pre-perturbation), and found 268/345 perturbation units to be orientation selective (Hotelling’s T2-test, p<0.01). Interestingly, the preferred perturbation orientation often matched the preferred orientation of neurons (Fig. 2d, e). Neurons with significant orientation selectivity showed a perturbation response tuning similar to that obtained with the drifting gratings (Suppl. Fig. 3d). The distribution of the difference between the preferred visual and perturbation orientation was centred close to zero (0°±30° for n=104/268 neurons, Fig. 2e). We did not find a significant relationship with direction preference of the neurons (Suppl. Fig. 3c). These results suggest that the perturbation responses are more influenced by sensory feature selectivity of the neuron than by running direction.

The perturbation responses, while not consistent with trial-by-trial sensorimotor expectations, might be related to expectations built up by prior exposure to the visual stimuli. To test this, we recorded from two groups of animals: one group (n=7 mice, 37 recording sessions) was exposed to the drifting gratings (without perturbations) over 8-13 days, while the other group (n=3 mice, 10 recording sessions) did not experience any stimuli before the first recording session. We found a similar percentage of perturbation units in both groups (60% for naïve and 58% for experienced mice, Suppl. Fig. 2), suggesting that prior exposure did not influence the occurrence of perturbation responses.

What properties of a neuron determine whether or not it is perturbation responsive? While, expectation was a potential hypothesis, our evidence did not support this, and instead found a larger role for stimulus orientation. We therefore investigated whether additional stimulus properties predict neural responses to visual flow perturbations. Given that the perturbation responses occurred when the drifting grating was suddenly stopped (or made static), we hypothesized that the positively modulated perturbation units preferred static gratings over drifting gratings. To test this, we tested a range of speeds of the grating (different temporal frequencies at a fixed spatial frequency), moving along the naso-temporal (front-to-back) direction (Suppl. Fig. 4a; 3 animals, n=109 neurons, temporal frequencies tested: 0:0.5:8.5 cycles/s, Suppl. Fig. 4b,c). The speed preferences of perturbation units were consistent with our hypothesis (Suppl. Fig. 4h): positively modulated units mostly preferred static gratings whereas negatively modulated units mostly preferred speeds greater or equal than 3 cycles/s (MI>0, n=62; MI<0, n=47, Mann-Whitney U test, p = 1.3e-6, Fig. 3c). We then considered the possibility that perturbation responses could be explained by direction selectivity. For instance, a grating moving in the opposite or orthogonal direction from that preferred by the neuron might suppress its firing rate. However, we found that the preference for slow speeds was not explained by a preference for other motion directions (Suppl. Fig. 4i). These data suggest that a preference for static or slower moving gratings explains the increases in firing rate observed in response to the visual flow perturbation, irrespective of predictions based on running direction.

**Fig. 3:**
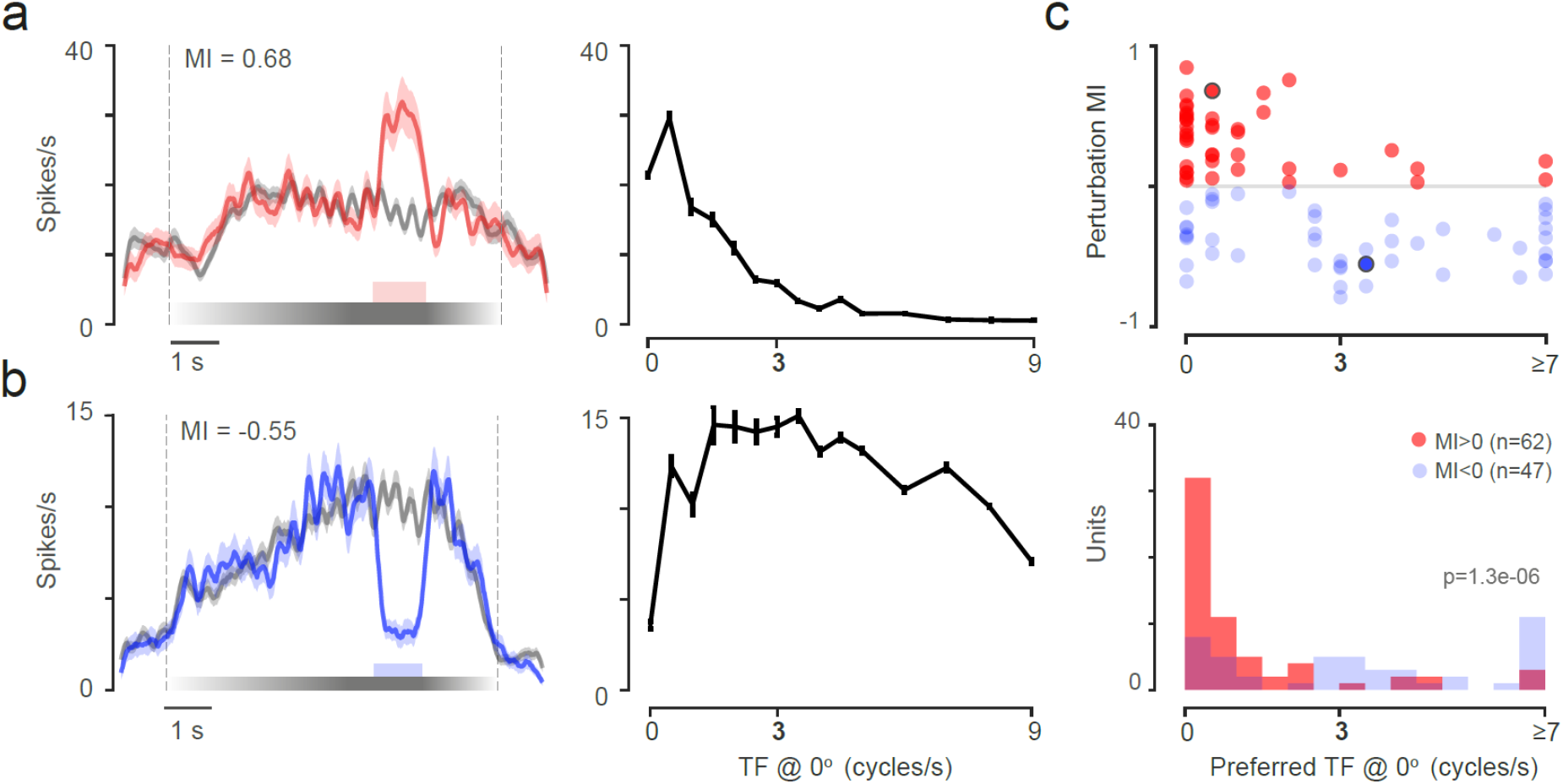
Temporal frequency tuning can explain perturbation responses. **a,** Example neuron positively modulated by the visual perturbation. Left: temporal frequency tuning. Right: mean response to perturbation trials (red trace) and non-perturbation trials (black trace). Shaded area indicate standard error of the mean, s.e.m. MI stands for modulation index of perturbation response. **b,** like a but with example neuron negatively modulated by the visual perturbation. **c,** Top: perturbation modulation index as a function of the preferred speed for all units. Circled data points are units shown on **a** and **b**. Bottom: Distribution of preferred speed for positively (red, n=62) and negatively (blue, n=47) modulated perturbation responsive units. All preferred speeds greater or equal to 7 cycles/s are grouped together for illustration purposes.

## Discussion

We have shown that a significant proportion of neurons of the visual cortex of mice respond to visual flow perturbations during passive viewing of drifting gratings. Although these responses were enhanced when animals were running, the presence or strength of perturbation responses did not depend on the direction of the previously experienced visual flow. Visual perturbation responses were instead better explained by the preferred orientation of the neuron, and preferences for static or slowly moving stimuli. This suggests that perturbation responses can be explained by visual feature tuning that is enhanced by locomotion, and do not require internal monitoring of prediction error relative to self-generated motion.

Neurons responding to visual perturbations are qualitatively similar to sensorimotor mismatch neurons previously described. Specifically, neurons have been classed as sensorimotor mismatch neurons when their responses match the following conditions^10^: they increase their activity in response to sudden stops of visual flow, and the response magnitude is at least twice as strong when animals run. We have shown here that many visual perturbation neurons can also satisfy these conditions. Using the same criterion, we found 13% of recorded neurons could be classed as sensorimotor mismatch neurons, which is in range of the percentages previously reported^10,13,23,24^ (which vary between studies from 5% to 39%). Our observation that visual perturbation units would be classed as sensorimotor mismatch neurons does not exclude the possibility that there are neurons that are indeed selective to true sensorimotor mismatch, especially in animals that have had extended experience of closed-loop conditions^13^. However, to assess the prevalence of purely sensorimotor mismatch neurons, one would have to discount effects of stimulus properties that explain visual perturbation sensitive units shown here. True visuomotor mismatch neurons should have greater responses to visual perturbations in closed-loop than during playback of the same stimuli while the animals are running, and these responses should be higher in the direction of expected visual flow during locomotion.

The visual tuning properties of neurons that respond to sudden stops (perturbation or mismatch) are consistent with typically reported properties of mouse visual cortex. The direction tuning of perturbation responses was not biased in the expected visual flow direction – the front-to-back direction. Instead, we found perturbation responses to be better predicted by the orientation tuning of the neurons. Previous work characterising visual responses to sudden stops in running animals have only explored responses to front-to-back motion. Neurons responding positively to a sudden stop also prefer slow speeds. That is, they prefer the speed experienced during the sudden stop, rather than that experienced during the visual flow. This simple visual tuning is enhanced during running, a well-known property of visual neurons^17–19^.

For a neural network to implement predictive coding, a fraction of neurons need to encode prediction errors^11^. In the case of visual flow, high prediction error would be represented by high activity levels when an animal is running fast, but experienced visual flow is slower. The sensorimotor mismatch model suggested for encoding prediction errors in V1 combines an excitatory efference signal (motor command) and an inhibitory sensory feedback^12,13^. However, our data show that it is possible that a preference for low visual flow and high running speeds, both potentially excitatory inputs, can produce responses consistent with a prediction error signal. Our observations also explain why the distribution of visual speed tuning preferences across the layers of V1 are consistent with the prevalence of neurons responsive to sensorimotor mismatch^12^: superficial layers that have a larger fraction of mismatch neurons compared to deeper layers, also have a larger fraction of neurons preferring low temporal frequencies (based on the Allen Institute data set^25^; Suppl. Fig. 5). Therefore, encoding of prediction error in the visual cortex can be achieved by the convergence of distributed visual tuning properties with modulation of responses by running.

Features consistent with the predictive coding hypothesis have been observed in primary sensory areas in responses to the omission of expected stimuli^26,27^ and suppression of responses to expected sounds^9^. Similarly, internal representations of spatial location predict the responses of upcoming stimuli^26^ and modulate the responses to identical sensory stimuli^28,29^. These responses are often explained as the difference between efference information and the sensory signal, but our results propose a simpler model that may be useful: the convergence of distributed sensory and motor response codes. Incorporating and accounting for these factors provides a richer testbed for theories of sensory coding during self-generated actions.

## Acknowledgements

The authors declare no conflict of interest. This work was supported by The Sir Henry Dale Fellowship from the Wellcome Trust and Royal Society (200501); and by the Human Frontier in Science Program (RGY0076/2018). We thank Sylvia Schröder, Bilal Haider and Samuel Solomon for comments on the manuscript.

## Supplementary Figures

**Suppl. Fig. 1.**
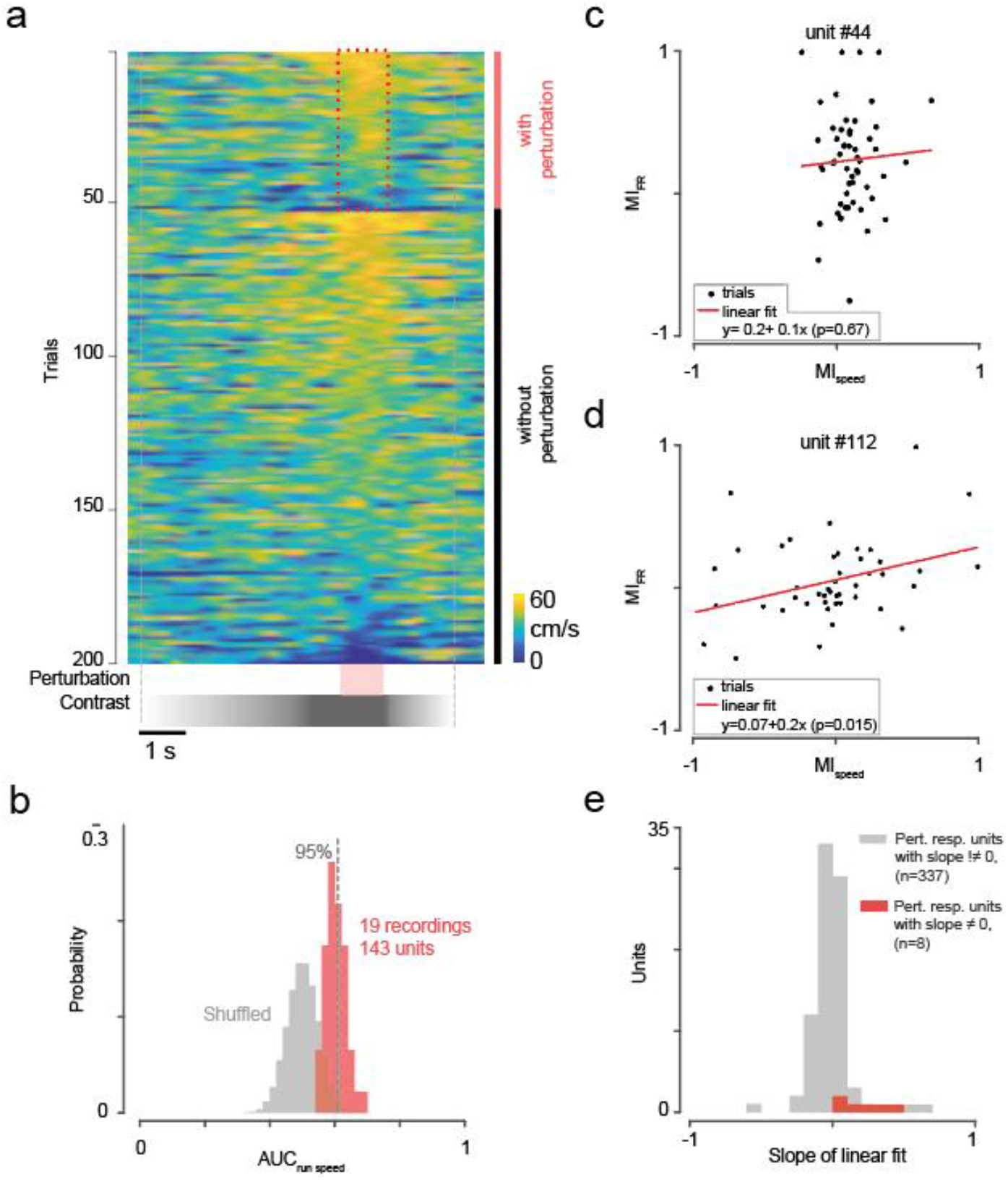
Visual perturbation responses are not explained by perturbation-induced changes in running behaviour. **a**, Top: normalised running speed during every trial of an example session. Bottom: contrast and visual flow perturbation period (TF = 3 → 0 Hz). **b**, Probability distribution of the area under the receiving operating characteristic curve of the logistic classifier trained with shuffled and actual running speed data. **c**, **d**, scatter plot of modulation indexes for firing rate (MI_FR_) and running speed (MI_speed_) for two example units. Modulation is computed as the difference between the mean firing rate (or speed) during the perturbation and the pre-perturbation period, and normalised by the second term. The linear fit (red line) measures the correlation between the two modulations. The *p*-value indicates the *t*-statistic of the hypothesis test that the slope is significantly different from zero or not. **e**, Distribution of the slope of the linear fits for all positively modulated perturbation responsive units. Significance is measured at the 1% significance level

**Suppl. Fig. 2:**
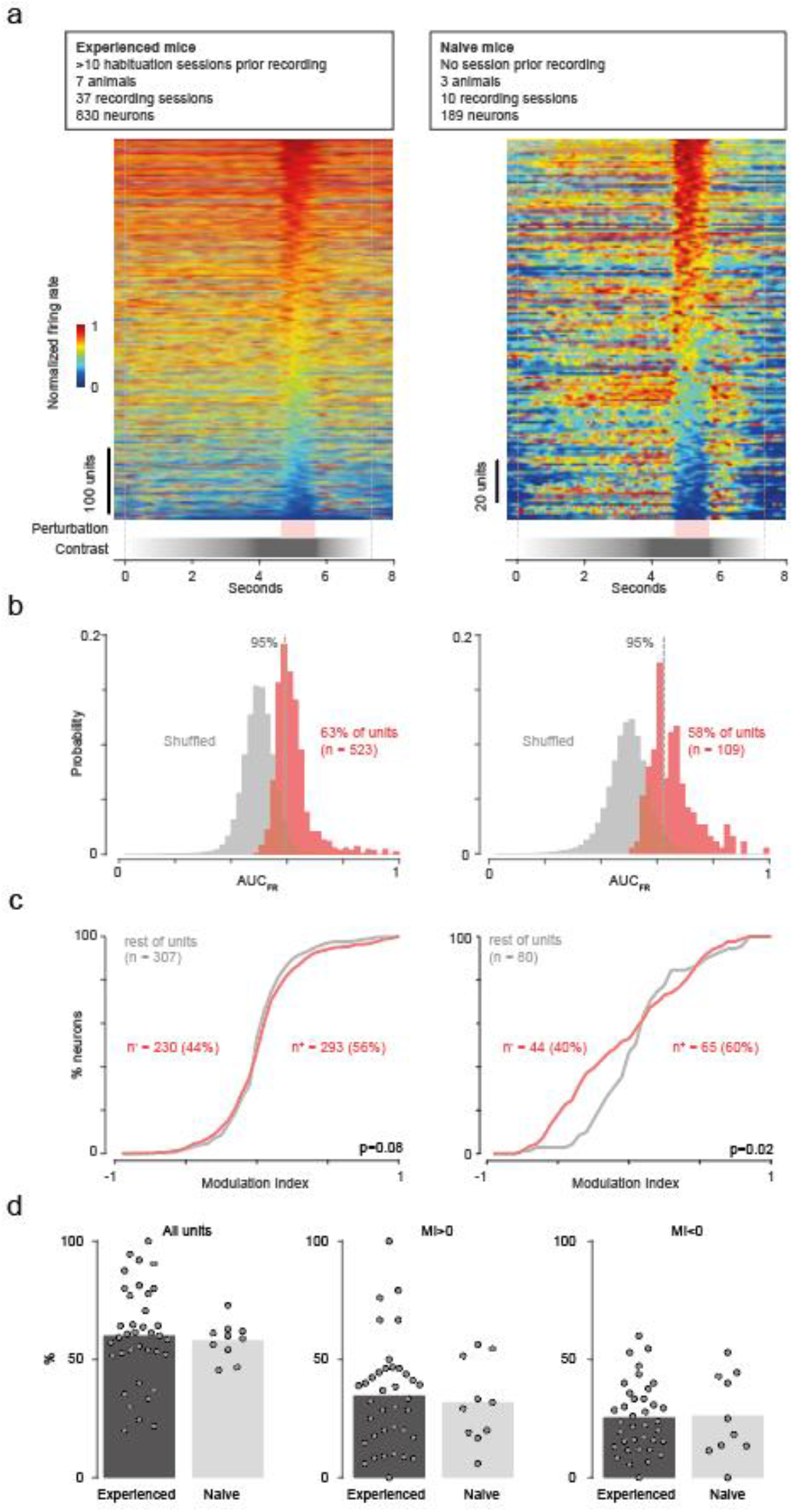
Perturbation responses are also present in naïve animals. **a**, Top: normalised mean response of all recorded units (n=189) for trials with perturbation. Bottom: and visual flow perturbation period (TF = 3 → 0 Hz). Left: Experienced mice with at least 8 habituation sessions. Right: Naïve mice with no experience of visual stimulation. Same order applies to other panels. **b**, Probability distribution of the area under the receiving operating characteristic curve of the logistic classifier trained with shuffled and recorded neural data. **c,** Probability cumulative distribution of the modulation indexes (MI) of perturbation responsive units (red) and the other units (gray). P value for two-sample Kolmogorov-Smirnov test.. **d,** Breakout of percentages of perturbation responsive units per session for experienced (n=37) and naïve mice (n = 10).

**Suppl. Fig. 3:**
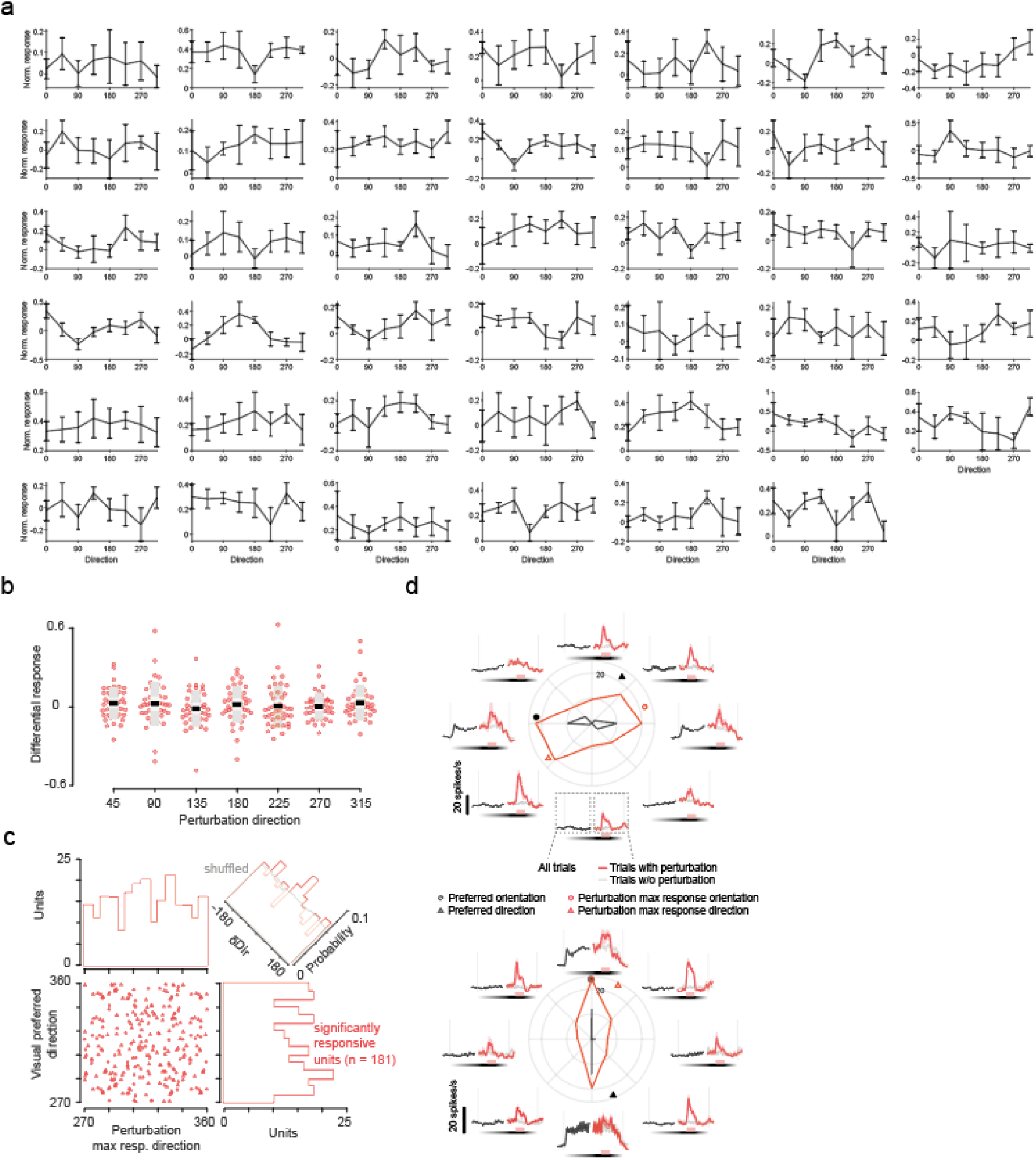
Population responses are not affected by visual flow direction. **a,** Normalised perturbation responses of individual session populations (n>1) for all directions tested (0:45:315°). **b,** Perturbation responses for all neuron ensembles of units recorded in a session (n = 41, number of units per session ranges between 2 and 24). Colours of box plots indicate mean (black), s.e.m. (light gray), s.t.d. (dark grey). **c**, Scatter plot of preferred direction angle for grating stimulus and visual perturbation. Only units tuned to any grating direction are shown (Hotelling’s t-squared test, p<=0.01 n = 268). **d**, Mean responses of two example units for the different grating directions.

**Suppl. Fig. 4:**
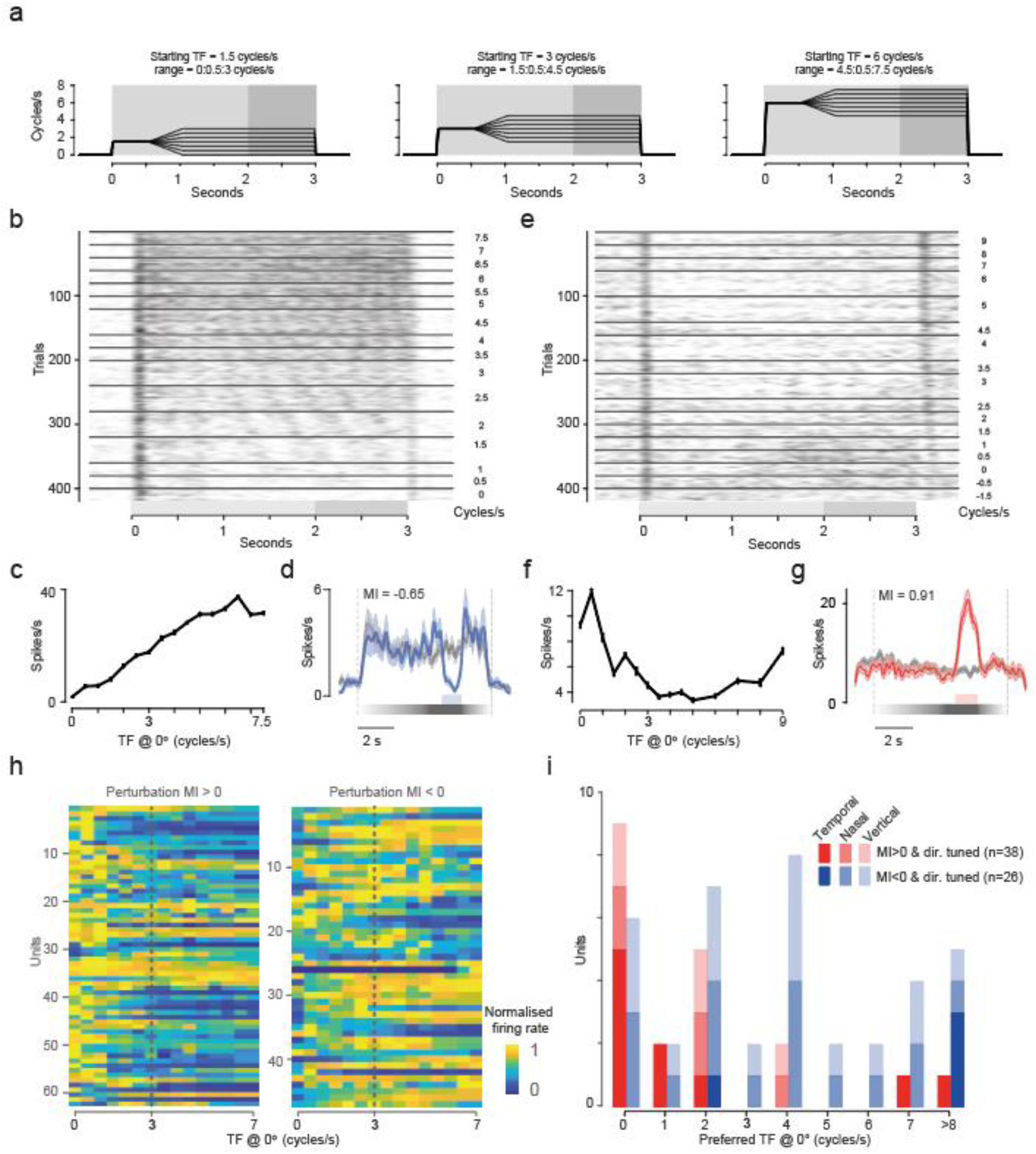
Temporal frequency tuning of perturbation responsive units. **a,** Temporal frequency profiles shown during trial (0 → 3s, grey area). We used 3 starting TF’s: 1.5, 3, and 6 cycles/s. After 0.5 s, this was increased or decreased at a random rate −3:1:3 cycles/s^2^ for 0.5 seconds and then kept constant (see black traces). To compensate for onset transient responses, the temporal frequency tuning was evaluated in the last second of the trial (dark grey area). **b, c, d** example neuron negatively modulated by the visual perturbation: **c** is the tuning curve of spiking activity during all trials for all temporal frequencies tested;. Mean response to perturbation (red trace) and non-perturbation trials (black trace) is shown in d. Shaded area indicate standard error of the mean, s.e.m. **e, f, g,** like b, c, d but with example neuron positively modulated by the visual perturbation. Please not that in this session, the temporal frequency changed for 1 second instead of 0.5 second, like shown in a. Hence TF values reached higher (and lower) values. We only considered TF’s ≥ 0. **h,** Heat map of speed tuning curves computed along the temporal direction for the perturbation responsive units. Dotted line indicates the temporal frequency used in the main stimulus (3 cycles/s) before the perturbation. Left heat map shows larger blue area for speeds >3 cycles/s compared to right indicating a preference for low temporal frequencies of positively modulated perturbation responsive units. **i**. Distribution of preferred temporal frequencies for neurons responsive to visual perturbation (MI>0 in red and MI<0 in blue) and significantly direction tuned (direction dot product test p<0.01). Preferred directions are grouped according to the closest temporal, nasal, and vertical (superior and inferior) axes (angle difference <45°).

**Suppl. Fig. 5:**
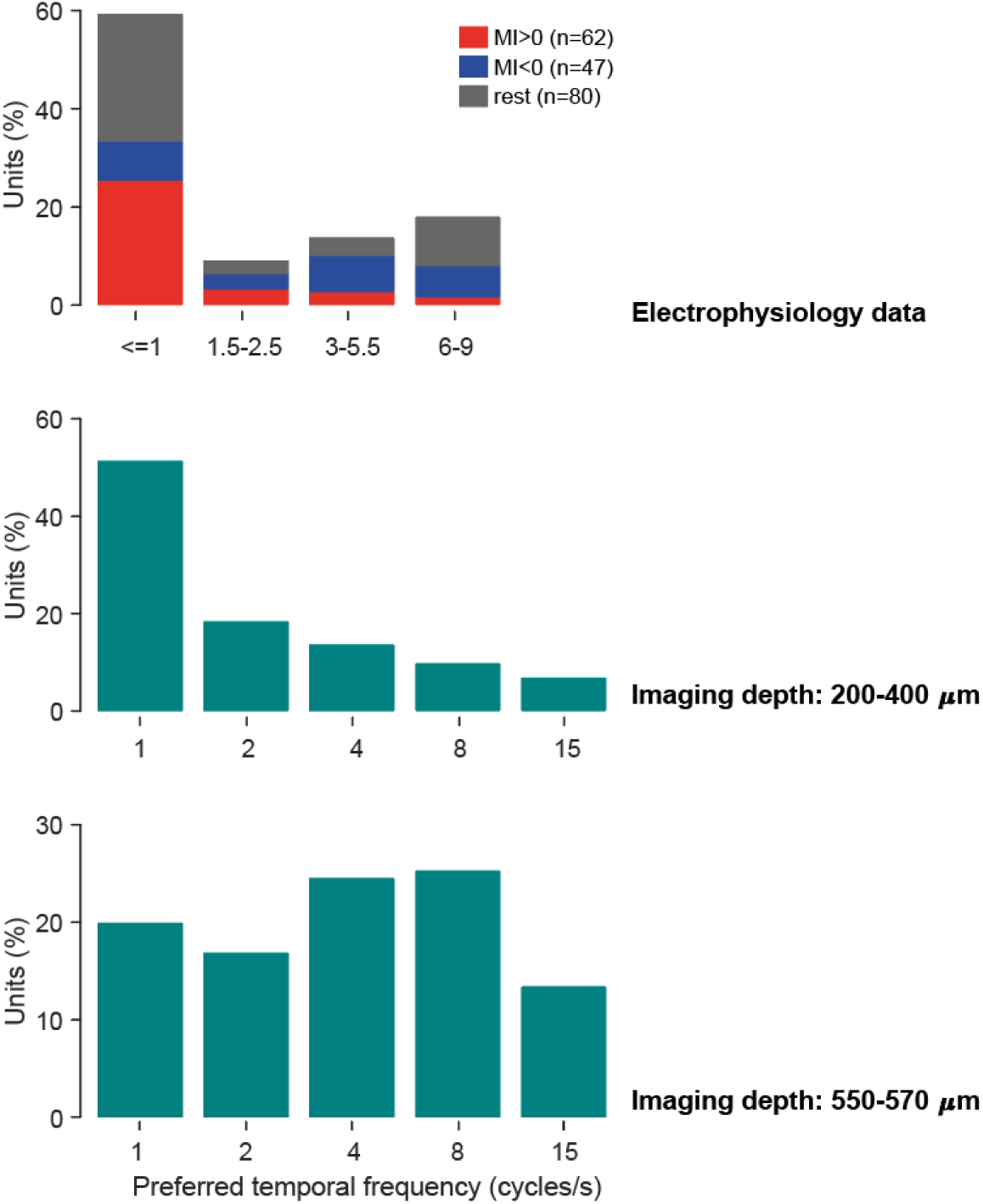
Comparison of preferred temporal frequencies with data from Allen Brain Institute. **a,** Distribution of the preferred temporal frequencies for neurons of our dataset (top), Allen Brain Institute for imaging depths of the mouse visual cortex between 200 and 400 μm (middle, n = 9059), and for depths of 500-570 μm (bottom, n = 261). Data retrieved from the online data portal (url: http://observatory.brain-map.org/visualcoding). Neurons shown here are significantly responsive to drifting grating (p<0.01, n = 9059/15619 for superficial layers and n = 261/573 for deeper imaging depths.)

## Methods

### Surgery

All experiments were performed in accordance with the Animals (Scientific Procedures) Act 1986 (United Kingdom) and Home Office (United Kingdom) approved project and personal licenses. The experiments were approved by the University College London Animal Welfare Ethical Review Board under Project License 70/8637. The mice (n = 10 C57BL6 wild-type, 7 females and 3 males, age 16-24 weeks) were housed in groups of maximum five under a 12-hour light/dark cycle, with free access to food and water. All electrophysiological recordings were carried out during the dark phase of the cycle.

Mice were implanted with a custom-built stainless-steel metal plate on the skull under isoflurane anaesthesia. The area above the left visual cortex was kept accessible for electrophysiological recordings. Seven days following the surgery mice underwent the first habituation session. Following the habitation period (one session per day, 8-13 days), a craniotomy was performed over V1, centred at 2 mm lateral to sagittal midline and 0.5 mm anterior to lambda). We dura was left intact to preserve the brain tissue, and maximise recording time. Mice were allowed to recover for 4-24 hours before the first recording session. Multiple recording sessions were performed on each animal (one per day, n = 37 recordings, min 2, max 9). Three animals (naïve) were exposed to only grey screen during the habituation sessions (5-10 sessions) and were exposed to the visual stimulus for the first time on the day of the recording sessions.

### Visual Stimulus

The display apparatus was similar to those used in previous studies^30,31^. Mice were head-fixed on a polystyrene wheel^28^ (radius 10 cm), with their heads positioned in the geometric centre of a truncated spherical screen onto which we projected the visual stimulus. The visual stimulus was centred at +60° azimuth and +30° elevation and had a span of 120° azimuth and 120° elevation.

The visual stimulus was designed using BonVision^32^, an open-source visual environment generator based on graphical programming language of the Bonsai framework^33^. A session was structured in trials during which a drifting sinusoidal grating was presented for approximately 7.3 seconds (sinusoidal grating with spatial frequency of 0.04 cycles/° and temporal frequency of 3 cycles/s). The grating was made visible by increasing the contrast from 0 to 0.8 in the first 240 frames of the trial. Contrast remained at 0.8 for the following 100 frames and then decreased to zero in 100 frames. Total trial duration was 440 frames at 60 Hz frame rate, i.e. 7.3 seconds. The direction of the grating was randomly picked between eight directions: 0°:45°:315°. Each grating direction was shown at least twenty times. For each combination of directions a trial was shown with zero contrast. The inter-trial interval varied randomly between 1 to 2.5 seconds. The firing rate during this period and the zero-contrast trial was used as a baseline. Habituation sessions for naïve mice were run with a gray screen.

The visual perturbation was presented in a random 25% of the trials, during the recording sessions. The perturbation onset happened between 30 and 40 frames after the contrast had reached its highest value (0.8), and the perturbation lasted between 60 and 70 frames.

### Data Analysis

#### Spike sorting and clustering

To record the neural activity we used multi electrode array silicon probes with two shanks and 32 channels (ASSY-37 E-1, Cambridge Neurotech Ltd, Cambridge, UK). Electrophysiology data was acquired using an OpenEphys acquisition board^34^.

The electrophysiological data from each session were processed using Kilosort version 1 (Pachitariu et al., 2016). Spike times were synchronised with the behavioural data by aligning the signal of a photodiode that detected the visual stimuli transitions (PDA25K2, Thorlabs, Inc., USA). They were then analysed conjointly in Matlab R2019a. Firing rate was sampled at 60 Hz and smoothed with a 300-ms Gaussian filter.

#### Logistic regression classifier

To estimate the reliability of the responses to the visual perturbation we trained a trial classifier based on a logistic regression model (Matlab function *fitglm*). We used six parameters from each trial as inputs for the model:

1. the ratio of mean firing rate during the perturbation period (*R_pert_*) and during the second preceding it (*R_pre–pert_*):

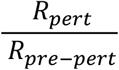
2. the difference between them:

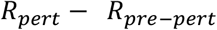
3. their sum across the whole period:

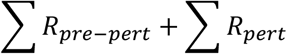
4. the summed firing rates during the perturbation period:

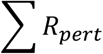
5. a depth of modulation index:

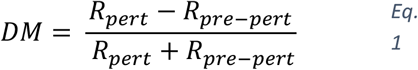
6. a response modulation index:

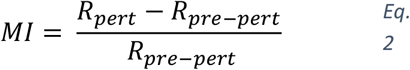

The output of the classifier was the identity of the trial: 1 for perturbation or 0 for non-perturbation trial. We then computed the receiving operating characteristic curve (true positive rate VS negative positive rate) and its area under this curve (AUC) was used as reliability parameter. We then shuffled the trial type 1000 times and rerun the above steps to evaluate the statistical significance of this metric. If the AUC of each unit was greater than 95% (p < 0.05) of the AUC’s computed from all shuffled data, we considered the unit as perturbation responsive.

#### Modulation index

We then quantified the magnitude of the perturbation response and identified whether a unit increased (MI>0) or decreased (MI<0) its firing rate during the perturbation, using the modulation index as shown in Eq. 2.

#### Responses during running

To evaluate the effects of running on perturbation responses we divided the perturbation trials in two groups based on the running state of the mice. If the mouse ran at speeds greater than 2 cm/s during the perturbation period and the second preceding it, we consider it a running trial. Other trials are considered stationary trials. As the animals were free to run or stand still as they wished, we often found an imbalance in the number of stationary and running trials. To better compare the two conditions, we selected only sessions that had at least 4 trials per condition. This allowed us to compare the activity of 174 neurons from 22 sessions. We measured the modulation of the running and stationary trials as the area under the response curve during the perturbation period (analytically is the sum of all firing rates) minus the area under curve evaluated during the pre-perturbation (1 second).

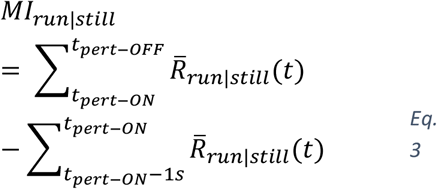

where 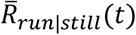 is the average of the mean instantaneous firing rates of either the run or still perturbation trials. To compensate for biases due to smaller number of trials for either condition, we compared the activity of 20 trials per condition by re-sampling 1000 times with replacement. An individual neuron was classified as significantly modulated by running if both the peaks and the mean responses computed for running trials *R_run_*(*t*) were larger than those computed for still trials *R_still_*(*t*) in 950/1000 of the cases (5% significance level).

#### Orientation and direction tuning

We estimated the mean responses to the drifting grating at different angles by measuring the mean firing rate in the first 3.5 seconds from the onset of the stimulus and subtracting the pre-stimulus mean firing rate. Similarly, the neural response for different perturbation directions were computed as the mean firing rate during such period, minus the pre-stimulus (inter-trial period) mean firing rate. We then estimated the preferred orientation and direction tuning of the neurons for the drifting grating using standard methods^35^. Briefly, responses to various directions and orientations were evaluated in either a direction or orientation vector space, and we evaluated the normalised length of the vector sum as a measure for direction and orientation selectivity:

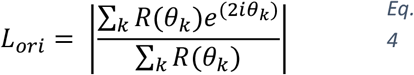

where *R*(*θ_k_*) is the response to angle *θ_k_*.

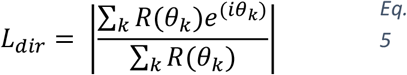

The preferred orientation or direction of a unit were estimated from the angles of the vectors in the numerator of Eq. 4 and Eq. 5, respectively. The significance of orientation tuning was computed using Hotelling’s *T*^2^-test, which is a multivariate generalization of the Student’s *T*-test. The significance of direction tuning was computed using “direction dot product” test. Significance level was set at 99%. For perturbation responses, we used the direction/orientation with the maximal responses, as the number of trials for each direction was too small to implement the aforementioned method.

#### Temporal frequency tuning

To estimate the temporal frequency tuning we used stimuli with similar specifications to those used for the visual perturbation stimuli (sinusoidal grating only at 0° direction and SF = 0.04 cycles/°, contrast = 0.8, Suppl. Fig. 4). This was presented only to naïve mice. Each trial commenced with a grating moving at either 1.5, 3, or 6 cycles/s. After one second (60 frames) there was a period of acceleration, where the temporal frequency linearly increased or decreased at a random rate between seven possible values (−3:1:3 cycles/s^2^) for 0.5 or 1 seconds (30 or 60 frames). At the end of the acceleration periods the grating speed was kept constant for 1.5-2 seconds (90-120 frames) followed by a grey inter-trial interval of 1 to 2.5 seconds. Twenty-one combinations of initial temporal frequencies and acceleration rates were presented twenty times each. The mean firing rates of the final second of all trials with a specific temporal frequency were used to calculate the speed tuning curve for the perturbation responsive neurons (n=109). Preferred temporal frequency was the temporal frequency with the maximal response.

